# Residual Complex I activity supports glutamate catabolism and mtSLP via canonical Krebs cycle activity during acute anoxia without OXPHOS

**DOI:** 10.1101/2022.09.26.509156

**Authors:** Dora Ravasz, David Bui, Sara Nazarian, Gergely Pallag, Noemi Karnok, Jennie Roberts, Daniel A Tennant, Bennett Greenwood, Alex Kitayev, Collin Hill, Timea Komlódi, Carolina Doerrier, Erich Gnaiger, Michael A Kiebish, Alexandra Raska, Krasimir Kolev, Bence Czumbel, Niven R Narain, Thomas N Seyfried, Christos Chinopoulos

## Abstract

Anoxia halts oxidative phosphorylation (OXPHOS) causing an accumulation of reduced compounds in mitochondrial matrix which impedes dehydrogenases. By simultaneously measuring oxygen concentration, NADH autofluorescence, mitochondrial membrane potential and ubiquinone reduction extent *in organello* in real-time, we show that Complex I utilized endogenous quinones to oxidize NADH under acute anoxia. Untargeted or [U-^13^C]glutamate-targeted metabolomic analysis of matrix and effluxed metabolites extracted during anoxia in the presence or absence of site-specific inhibitors of the electron transfer system inferred that NAD^+^ regenerated by Complex I is reduced by the 2-oxoglutarate dehydrogenase complex yielding succinyl-CoA supporting mitochondrial substrate-level phosphorylation (mtSLP), releasing succinate. Yet, targeted metabolomic analysis using [U-^13^C]malate also revealed concomitant succinate dehydrogenase reversal during anoxia yielding succinate by reducing fumarate, albeit to a small extent. Our results highlight the importance of quinone availability to Complex I oxidizing NADH, thus maintaining glutamate catabolism and mtSLP in the absence of OXPHOS.

## Introduction

Within the mitochondrial matrix of humans and the mouse — the laboratory animal model used in this study — NADH can be irreversibly oxidized in 16 and reversibly in 30 reactions (supplementary table 1). Although some of them participate in specialized pathways (such as steroid metabolism or bile acid biosynthesis) and are not ubiquitous, all mammalian mitochondria harbor Complex I (CI, NADH:ubiquinone oxidoreductase, EC 1.6.5.3). CI catalyzes the oxidation of matrix NADH by ubiquinone (UQ) to produce NAD^+^ and ubiquinol (UQH_2_), which is coupled to the translocation of four H^+^ across the inner mitochondrial membrane and the transfer of electrons downstream to FeS clusters (Hirst, 2013). Mindful of the limitations quantifying NADH oxidation by CI (Birrell and Hirst, 2013) and the enhancement of NADH autofluorescence by mere binding to CI (Blinova et al., 2008), it is difficult to decipher the extent of contribution of CI in altering the mitochondrial NADH/NAD^+^ ratio. Nevertheless, it is a textbook definition that cessation of the electron transfer system (ETS) due to lack of oxygen, pharmacological inhibition or genetic ablation of respiratory components leads to an accumulation of reduced compounds in mitochondrial matrix, exactly because CI is not able to oxidize NADH. In turn, this increase in matrix NADH/NAD^+^ ratio is expected to impair the function of matrix dehydrogenases, preventing mitochondria from catabolizing substrates in the citric acid cycle (Xiao and Loscalzo, 2019).

Having said that, the catabolism of glutamine through oxidative decarboxylation of 2-oxoglutarate (oxoglutarate, Og, α-ketoglutarate) during hypoxia - in addition to its anabolism by reductive carboxylation-is firmly established (Zhang et al., 2018), (Mullen et al., 2014), (Seyfried et al., 2020). The question arises, what provides NAD^+^ to 2-oxoglutarate dehydrogenase complex (OgDHC) in the oxidative decarboxylation branch of glutaminolysis? Spinelli et al showed that in hypoxia (1 % O_2_) CI is still able to deposit electrons into the ETS (Spinelli et al., 2021). This process was driven by the reverse operation of succinate dehydrogenase (CII) reducing fumarate, supported by the high UQH_2_/UQ ratio. In their work (Spinelli *et al*., 2021), residual activity of CI in hypoxia was implied, something that has been previously proposed as plausible by mathematical modelling, even in anoxia (Chinopoulos, 2020). Mindful that Complex IV (CIV) exhibits a sufficiently high affinity for O_2_ that could maintain partial activity even in 1 % O_2_ (Gnaiger et al., 2000), Spinelli et al generated cells lacking key components of either CIV or Complex III (CIII) rendering them incapable of using O_2_ in the ETS (Spinelli *et al*., 2021). However, the cell lines were constitutive knock-outs for these components and most variables were recorded after several hours, thus the effect of an acute hypoxia or anoxia could not be addressed. Investigating the effects of anoxia in the acute phase bear equal -if not greater-pathophysiological relevance, as any tissue experiencing lack of oxygen may quickly succumb. Here we used isolated mitochondria, free from confounding factors stemming from extramitochondrial processes and performed experiments during acute anoxic phase. Using a battery of untargeted and targeted metabolomic analysis, biochemical, electrochemical and fluorescence assays, we report that in isolated mitochondria experiencing acute anoxia, the residual activity of CI is sufficient to provide NAD^+^ to OgDHC and maintain the oxidative decarboxylation branch of 2-oxoglutarate supporting mtSLP. We further show that concomitant to this, CII also operates in reverse - albeit to a minor extent -, reducing fumarate. The critical factor for allowing these phenomena to unfold is the endogenous pool of mitochondrial quinones.

## Results

### Complex I remains partially active during acute anoxia

Mindful that CI oxidizes one molecule of NADH to NAD^+^ by reducing UQ to UQH_2_ in an equimolar manner while pumping four protons out of the matrix as shown in figure 1A, Jin and Bethke derived a model for CI activity using non-equilibrium thermodynamics (Jin and Bethke, 2002). We used their rate equation to generate a 3D plot of CI activity (expressed in nmol e^-^·min^-1^·mg^-1^) and input very wide ranges of UQH_2_/UQ and NAD^+^/NADH ratio reported in the literature (Turunen et al., 2004), (Galinier et al., 2004), (Yamamoto and Yamashita, 1997), (Kroger and Klingenberg, 1973), (Kulkarni and Brookes, 2019).

**Figure 1:**
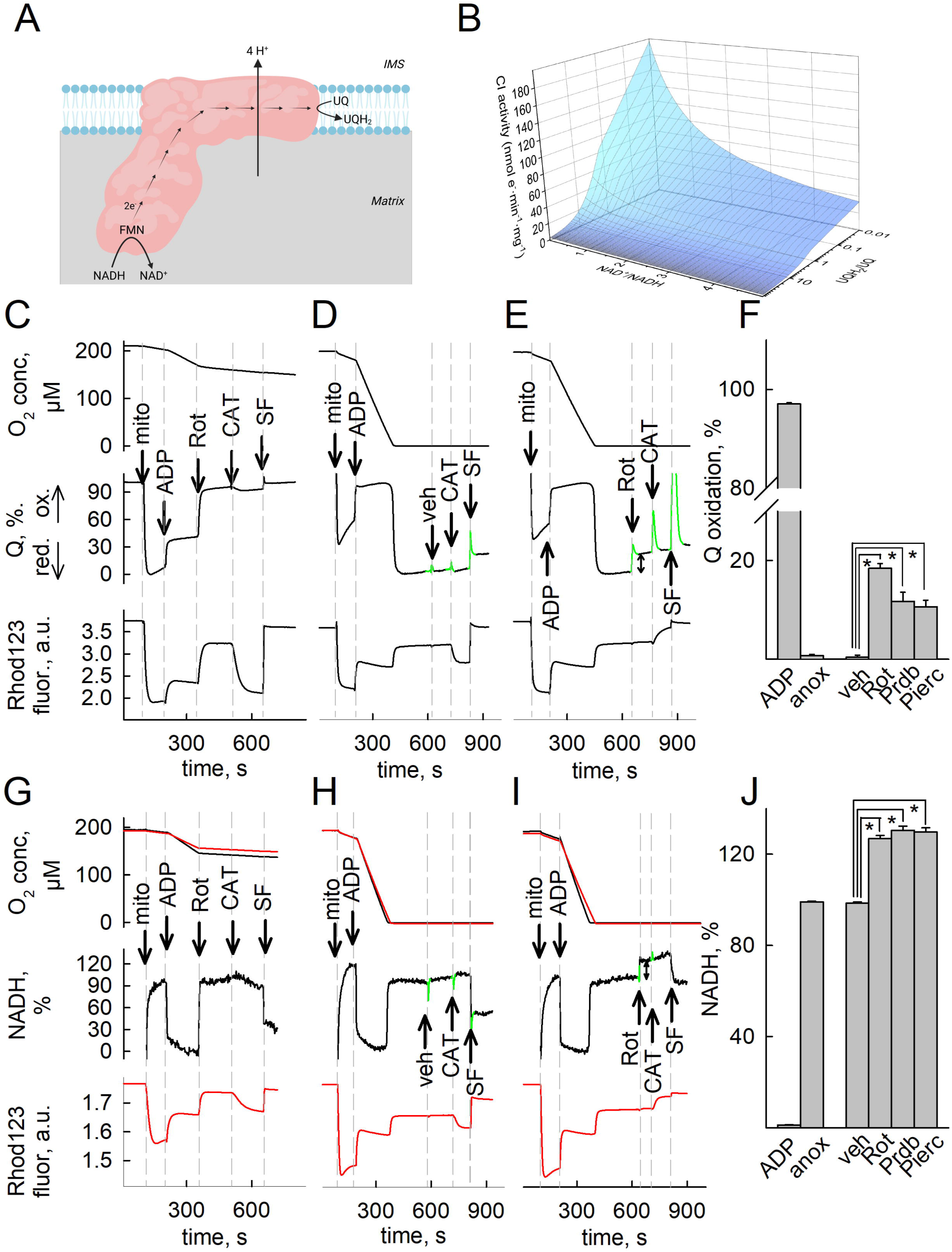
Complex I remains partially active during acute anoxia. A: Scheme illustrating the reaction catalyzed by CI (created with BioRender.com). B: 3D plot depicting CI activity as a function of UQH_2_/UQ, NAD^+^/NADH for Δ*Ψ*_mt_ −100 mV, pHin 7.35 and ΔpH=0.1; these values were taken from measurements in the present study and from those published in (Vajda et al., 2009). CI activity (J_CI) is the product of the rate equation formulated as J_C1 = Vmax * [NADH]/[NAD^+^]total pool * [Q]/[Q]total pool * F_T_, where F_T_ is thermodynamic drive. All concentrations are in matrix, [NAD]total pool = [NAD^+^] + [NADH]; [Q]total pool = [Q] + [QH_2_]. C, D, E, G, H, I: oxygen concentration (top panels, in μM) recorded simultaneously with either UQ redox state (C, D, E, middle panels) or NADH autofluorescence (G, H, I, middle panels) and/or rhodamine 123 fluorescence (Rhod123, bottom panels) in isolated mouse liver mitochondria. Rhodamine 123 fluorescence indicative of Δ*Ψ*_mt_ (arbitrary units, a.u.) was recorded separately from NADH autofluorescence due to spectral overlap. Note that in G, H, I the presence of rhodamine 123 decreased OXPHOS respiration rate (compare black with red traces), thus anoxia commenced at slightly different times. This is similar to what has been reported for safranine O (Valle et al., 1986), (Krumschnabel *et al*., 2014). Mitochondria (mito), ADP (2 mM), vehicle (ethanol) or rotenone (rot, 1 μM), carboxyatractyloside (CAT, 1 μM), SF (SF6847, 0.25 μM) were added where indicated. Substrates were glutamate and malate (5 mM each) present in the buffer prior to addition of mitochondria. Panels are aligned in the x-axis. F, J: quantification of the ADP, anoxia, vehicle or CI-inhibitor induced changes in UQ (panel F) or NADH (panel J) signal, as illustrated by the bidirectional arrows in panels E and I, respectively. The peaks colored in green in panels D, E, H and I are artefactual caused by the additions of the drugs and are removed from all further quantifications. * p<0.05.

As shown in figure 1B, when UQH_2_/UQ is very high (>10) and NAD^+^/NADH very low (< 1) mimicking anoxic conditions, CI activity is predicted to retain < 20 % of its theoretical maximum. We therefore set to investigate i) if CI activity can indeed be experimentally demonstrated under anoxic conditions and ii) whether the fluxes of the reaction products, i.e. NAD^+^ and UQH_2_ are sufficiently high for maintaining downstream processes and - if yes - to what extent.

As a first step, we set up an assay to simultaneously monitor Q redox state, NADH autofluorescence, O_2_ concentration and membrane potential (Δ*Ψ*_mt_) during anoxia, in a (sub)second scale.We used isolated mouse liver mitochondria because of the high yield, purity and level of intactness achieved in this preparation as opposed to mitochondria obtained from brain or heart producing either a very low yield or a high fraction of broken mitochondria (Chinopoulos et al., 2009). [O_2_] was detected polarographically, Δ*Ψ*_mt_ fluorimetrically (inferred from rhodamine 123 preloading), NADH by its autofluorescence, and Q redox state electrochemically, the latter using coenzyme Q_2_ as a mediator between the mitochondrial UQ pool and the electrodes (Komlódi T, 2021). As shown in figures 1C-E [O_2_], UQ % and Δ*Ψ*_mt_ and in 1G-I [O_2_] and NADH % or Δ*Ψ*_mt_ were measured simultaneously in the same sample; panels are aligned in the x-axis. In the top panels [O_2_] is depicted, in the middle panels UQ % (C-E) or NADH % (G-I) and in the bottom panels rhodamine 123 fluorescence, indicative of Δ*Ψ*_mt_. Mitochondria were added where indicated, followed by ADP inducing OXPHOS respiration. The CI-specific inhibitor rotenone (or vehicle) was added before (figures 1C, G) or after (figures 1D, E, H, I) the chamber was depleted from O_2_ (marked as “anoxia”). Upon induction of anoxia the oxygen sensor detects no further changes in O_2_ concentration (figure 1D, E, H, I top panels; for detailed description of the sensor response during the transition from normoxic to anoxic conditions, see (Gnaiger, 2001)). Anoxia was associated with near simultaneous, precipitous reduction of UQ (figure 1D, E, middle panels) and increases in rhodamine 123 fluorescence (indicative of depolarization, figure 1D, E, H, I bottom panels) and abrupt increases in NADH autofluorescence (figures 1H, I middle panels). As expected, ADP and anoxia conferred an increase-*vs* a decrease in UQ oxidation levels (figures 1D, E, middle panels), respectively, and accordingly, a decrease-vs an increase in NADH levels (figure 1H, I middle panels). However, it is also obvious that rotenone inhibited UQ reduction to UQH_2_ whether O_2_ was present (figure 1C, middle panel) or not (figure 1E, middle panel). The peaks (pseudocolored in green in middle panels 1D, E, H, I) are artefacts produced by the addition of chemicals; from such experiments we pooled the values in the UQ signals conferred by the additions (as indicated by the bidirectional arrow in 1E, middle panel) and show them in bar graphs in figure panel 1F. Arbitrarily, we assigned the effect of ADP and anoxia as the 100 % and 0 % internal reference points of UQ oxidized state scale, respectively. Furthermore, addition of rotenone to anoxic mitochondria yielded a further increase in NADH fluorescence (figure 1I, middle panel). Pooled values in NADH signals conferred by the additions (indicated by the bidirectional arrow) are shown in bar graphs in figure panel 1J (NADH levels after the addition of ADP and commencement of anoxia were arbitrarily assigned as the 0 % and 100 % internal reference points of the scale, respectively). The rotenone-induced increases in UQ and NADH suggest that CI was operating in forward mode; although CI is fully reversible (Drose et al., 2016), (Kotlyar and Vinogradov, 1990) the conditions allowing reversibility *in organello* are extreme: by plotting CI forward-*vs* reverse operation as a function of UQH_2_/UQ, NAD^+^/NADH, matrix pH (pH_in_) and ΔpH across the inner mitochondrial membrane and assuming that anoxia clamps Δ*Ψ*_mt_ of isolated mouse liver mitochondria to ~-100 mV (Kiss et al., 2014), there can be only two circumstances upon which CI may operate in reverse: as shown in supplementary figure 1, CI operates in reverse if either i) UQH_2_/UQ >100 and NAD^+^/NADH =10 and pHin=7.35 and ΔpH is between 0.8-1.0 (panel D), or if ii) UQH_2_/UQ >100 and NAD^+^/NADH =10 and pHin= 8.35 and ΔpH is between 0.5-0.8 (panel E). Alternatively, these values could reach more realistic numbers if mitochondria became more polarized during anoxia (delineated by the dotted areas in the same panels), which on the other hand, is extremely unlikely. Overall, it is probably impossible to achieve such conditions in intact mitochondria. Finally, as shown in figures 1C and 1G (bottom panels) inhibition of the adenine nucleotide translocase (ANT) by carboxyatractyloside (CAT) in rotenone-inhibited, but not anoxic mitochondria leads to a gain in Δ*Ψ*_mt_, commensurate with our previous data showing that ANT remains in forward mode when CI is inhibited (Chinopoulos et al., 2010). Likewise, addition of CAT to anoxic mitochondria (in the absence of rotenone, figures 1D and H, bottom panels) also leads to a gain in Δ*Ψ*_mt_, commensurate to our data published before (Kiss *et al*., 2014). However, CAT led to a loss of Δ*Ψ*_mt_ when CI was inhibited in anoxic mitochondria figure 1E, I bottom panels), implying ANT reversal (Chinopoulos, 2011b). The observation that the ANT was operating in forward mode when oxygen was absent or CI was inhibited and in reverse mode when both oxygen was absent and rotenone was added hints that rotenone affects mtSLP, the primary determinant of ANT directionality when OXPHOS is inhibited (Chinopoulos, 2011a). Alternatively, rotenone could have an off-target effect anywhere on pathways converging towards mtSLP, dictating ANT directionality. However, the alternative CI inhibitors pyridaben (Prdb) and piericidin A (Pierc) reproduced the effect of rotenone during anoxia, see figure 1F. Accordingly, a further increase in NADH autofluorescence was observed when added during anoxia, see figure 1J. It is noteworthy that the effect of the inhibitors on the UQ signal was unequal (strength of effect, in descending order: rotenone > pyridaben > piericidin A), while all three exerted the same effect in increasing NADH autofluorescence when added on top of anoxia. This probably reflects that although rotenone, pyridaben and piericidin A share a common binding domain at- or in the vicinity of the UQ reduction site (Ino et al., 2003), the exact sites of action can be more than one in addition to being partially overlapping (Degli Esposti, 1998). Nonetheless, addition of Prdb or Pierc during anoxia led to a CAT-induced loss of Δ*Ψ*_mt_, implying ANT reversal and therefore absence of mtSLP (supplementary figures 2A, B, C and D, lower panels). The effects of pyridaben and piericidin A on UQ and NADH redox state were similar to those of rotenone (supplementary figures 2A, B, C and D, middle panels). The lack of effect on these (and all other inhibitors used in this study) on inducing the permeability transition pore during anoxia (detected by changes in light scatter of the organelles) is shown in supplementary figure 3A. The potential connection of CI activity to mtSLP is provision of NAD^+^ for OgDHC which yields succinyl-CoA, in turn supporting mtSLP (Kiss et al., 2013). Because the rotenone-induced changes in NADH fluorescence and UQ % in anoxia were smaller than those conferred in the presence of oxygen (compare middle panels of figure 1G with 1I), we deduced that CI activity is only partially active during anoxia. Since we were unable to calibrate the UQ and NADH signal, it is consequently we did not quantify CI activity *in organello* during anoxia.

### Endogenous UQ pool is a finite source supporting partial CI activity during acute anoxia

To address the pool of UQ supporting CI activity during anoxia we recorded the % change in the UQ signal as a function of time elapsed from commencement of anoxia until the addition of CI inhibitors. NADH availability could not have been a factor of finiteness as it is an excess during the anoxic insult. As shown in figure 2, rotenone (A-C) or pyridaben (D) or piericidin A (E) was the CI inhibitor when using either glutamate and malate, or oxoglutarate, or oxoglutarate and malate as fueling substrates. Representative experiments used to estimate UQ % values as a function of CI inhibitor and/or substrate(s) are shown in supplementary figure panels 2E-K. It is evident that the longer the time elapsed from commencement of anoxia until addition of CI inhibitors the smaller the % change in UQ signal. As also expected, CAT induced loss of Δ*Ψ*_mt_ in all conditions irrespective of inhibitor or substrate(s) present or time elapsed, implying ANT reversal (supplementary figure 2). The data imply that the reducable pool of UQ is finite and/or the entity oxidizing UQH_2_ back to UQ cannot keep up with the rate of UQ reduction by CI. To address this, we fed mice with a diet devoid of vitamin K_3_. Omission of vitamin K_3_/K_1_ from the diet has been reported to influence the mitochondrial Q/menaquinones pool in laboratory rodents respectively (Kolesova et al., 1988), (Thijssen and Drittij-Reijnders, 1994), while supplementation of patients suffering from CIII deficiency with vitamin K led to improvement of ^31^P NMR measurements signifying oxidative phosphorylation *in vivo* (Eleff et al., 1984); the beneficial effects of dietary supplementation with vitamin K_3_-together with other metabolites-in patients with mitochondrial disorders is reviewed in (Marriage et al., 2003). Mice - and control littermates fed with regular chow-were kept in vitamin K_3_-deficient diet over the course of approximately 3 weeks. After each week, prothrombin time (PT) was measured as an indicator of vitamin K status; the vitamin K hydroquinone is the cofactor essential for the γ-carboxylase present on the membrane of the endoplasmic reticulum in the liver to convert multiple glutamate residues to γ-carboxyglutamic acid residues present in vitamin K-dependent proteins including coagulation factors II (prothrombin), VII, IX, and X. When there is a deficiency of vitamin K, these factors are undercarboxylated and circulate in this form. The functional consequence of the lost posttranslational carboxylation is the reduced binding affinity of the undercarboxylated coagulation factors for calcium associated to phospholipid membranes resulting in inadequate activity of these factors and slower blood clotting. In human clinical practice the levels of functional vitamin K-dependent factors are routinely assessed with measurement of PT, the prolongation of which is one of the diagnostic criteria of bleeding related to vitamin K deficiency (Marder, 2012). Clinical data unambiguously support the value of prothrombin time as a marker of functional prothrombin concentration (Balendran et al., 2017), but this assay detects only gross changes in the vitamin K pool, thus, it is not a sensitive indicator of vitamin K status (Card et al., 2020). Severe vitamin K deficiency in the rat, produced by strictly nutritional means, does not impair the efficiency of oxidative phosphorylation in liver mitochondria, with NADH-linked substrates (Paolucci et al., 1963). As shown in figure 2F, PT was increased in mice kept in vitamin K3-deficient diet. In post-day 16 after implementing the vitamin K_3_-deficient diet total liver UQ levels were diminished, see figure 2G; however, by post-day 24 the trend was reversed, which means the dietary intervention did not affect UQ content. This discrepancy is also reflected in the results by Thijssen and Drittij-Reijnders showing a very large increase in the standard deviation of PT upon omitting vitamin K from the diet of rats over time (Thijssen and Drittij-Reijnders, 1994). UQ % upon addition of rotenone did not differ in anoxic mitochondria obtained from mice kept on vitamin K_3_-deficient diet for 3 weeks *vs* control littermates, see figure 2H. Although it is possible to use transgenic mice with constitutive ablations in genes coding for proteins participating in the ubiquinone synthesis pathway, such mitochondria would inherently exhibit severe OXPHOS deficiencies to the point that ANT directionality could not be addressed, since such assays require fully polarized mitochondria with very high respiratory control ratios (Chinopoulos, 2011a). Despite that we could not manipulate the pool of endogenous UQ by dietary means, the data above support the notion that UQ availability is critical for CI activity during anoxia. However, this does not exclude the possibility of UQH_2_ also being simultaneously oxidized by some other entity during anoxia.

**Figure 2:**
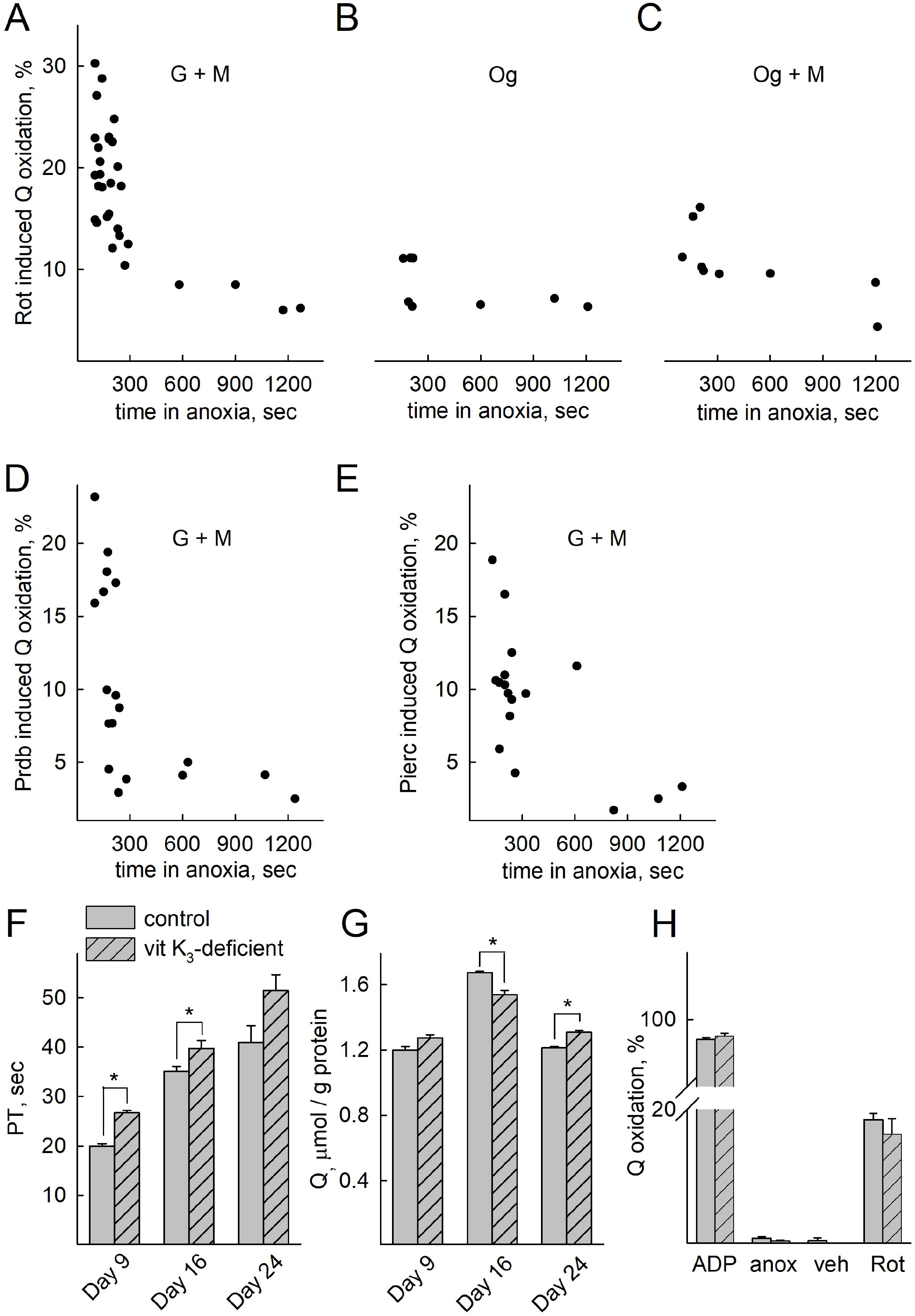
Endogenous UQ pools are a finite source supporting partial CI activity during acute anoxia. A-E: The CI inhibitor-induced change in UQ signal (% scale) plotted as a function of time elapsed from the onset of anoxia until the addition of CI inhibitors (time in anoxia, sec). For panels A, B, C, the CI inhibitor is rotenone (1 μM); for D, pyridaben (1 μM); for E, piericidin A (1 μM). Substrates are indicated in the panels (all at 5 mM). F: PT time in mice fed regular *vs* vitamin K3-deficient diet for 3 weeks; G: total quinones extracted from the livers of mice fed regular diet *vs*vitamin K3-deficient diet; H: quantification of the rotenone (Rot) induced changes in UQ signal from mitochondria obtained from mice fed regular-*vs* vitamin K3-deficient diet; veh: vehicle (for rotenone, i.e. ethanol). * p<0.05.

### CIII does not affect CI operation during anoxia to an appreciable extent

Having established that CI exhibits residual activity even in anoxia thus reducing UQ to UQH_2_ and mindful that the latter is the substrate of CIII (under normoxic or insufficiently hypoxic conditions), we tested the effect of CIII inhibitor myxothiazol on UQ and NADH redox levels. As shown in figure panel 3A, the presence of myxothiazol led to a minor though statistically significant decrease in UQH_2_ oxidation levels when added in anoxia, reflecting a minor CIII activity. This change was enhanced by addition of rotenone prior to myxothiazol (both added after anoxia), but UQH_2_ levels (figure 3B) were higher, implying that in anoxia, changes in UQ pool are mostly dictated by CI and not CIII. Rotenone and pyridaben (statistical significance was not reached with piericidin A) added after myxothiazol still implied residual CI activity even with CIII already inhibited. Similarly, the presence of myxothiazol before (figure 3C) or after (figure 3D) CI inhibitor(s) did not abolish NADH changes conferred by the CI inhibitor(s). The data argue that in anoxia there is a minor CIII activity, but most importantly CI residual activity is not solely influenced by inhibition of CIII. The electron acceptor(s) for this minor CIII activity were not sought further.

**Figure 3:**
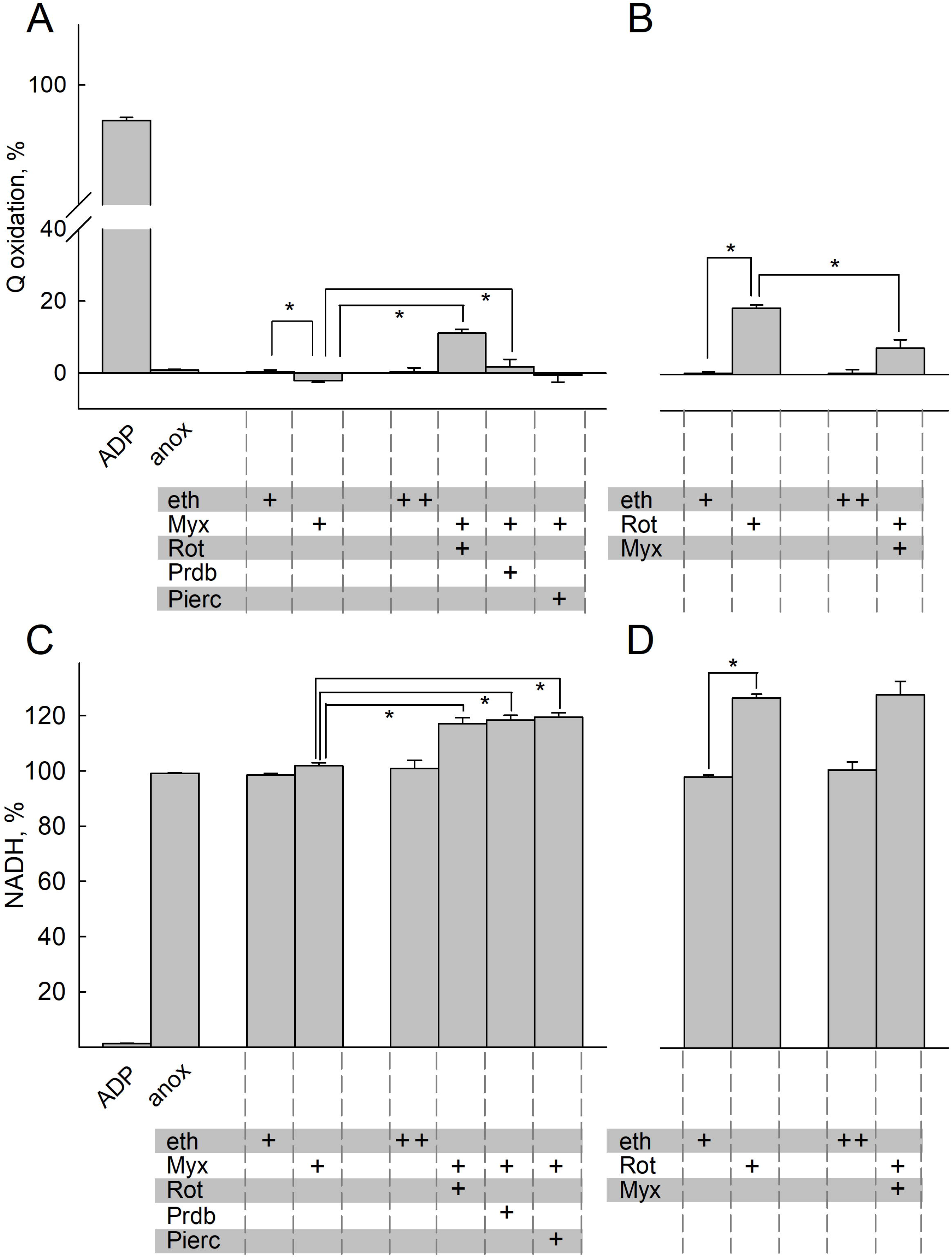
CIII does not mask CI operation during anoxia. Quantification of inhibitor-induced changes in UQ (A, B) or NADH (C, D) signal. Substrates and/or inhibitors present as indicated by the plus (+) signs at the bottom of the panels. Eth: ethanol; Myx: myxothiazol; Rot: rotenone; Prdb: pyridaben; Pierc: piericidin A. The additions of the compounds mentioned in the bottom of each panel were made from top to bottom. Data pooled from 3-37 independent experiments. * p<0.05. Whenever two crosses are indicated within a single box in the bottom of a panel mentioning additions of chemicals, they signify double addition of the vehicle.

### CII competes with CI for the same UQ pool during acute anoxia

Spinelli et al reported that in hypoxia or in cells genetically modified to lack CIII or CIV, the high UQH_2_/UQ ratio supported the reverse operation of CII, and as a consequence of this, CI remained partially operational (Spinelli *et al*., 2021). However, for the reasons reviewed in (Chinopoulos, 2019), the reverse operation of CII is unfavored, although not precluded. In the same line of thought, the group of Brookes reported that in cardiac ischemia, succinate accumulates mostly through canonical Krebs cycle activity through which the oxidative decarboxylation of 2-oxoglutarate produced from glutamine also takes place (Zhang *et al*., 2018). However, they did also report that CII was also operating in reverse, albeit to a moderate extent. The conundrum of whether CII operates in forward or reverse is relevant to the present study, because the reverse operation of CII yielding UQ would essentially be the means for maintaining CI operation forming UQH_2_, but in this case the finiteness of endogenous UQ pools would be rendered irrelevant. We therefore examined the effect of CII inhibitors atpenin A5 and malonate added before or after the addition of CI inhibitors during anoxia, and measured UQ oxidation % and NADH %. As shown in figure panel 4A, addition of any CI inhibitor after any CII inhibitor in mitochondria that were previously respiring on glutamate and malate (5 mM, each) led to a greater increase in UQ implying a greater inhibition of UQ reduction than addition of CI inhibitor alone (marked by a dash line), in addition to conferring an increase in NADH depicted in figure panels 4D; furthermore, addition of any CII inhibitor alone during anoxia did not lead to a statistically significant increase in UQH_2_/UQ compared to vehicle. However, addition of malonate (but not atpenin A5) during anoxia led to an increase in NADH autofluorescence, see (figure 4D). Addition of atpenin A5 after any CI inhibitor led to a greater increase in UQ, implying a greater inhibition of UQ reduction (figure 4B, marked by a dash line). Addition of atpenin A5 after any CI inhibitor did not lead to a statistically significant increase in NADH levels during anoxia, confirming that when CI is inhibited the contribution of CII to matrix NADH levels is irrelevant, see figure 4E. Note that while both CI and CII operate in the direction of UQH_2_ formation, this is not masked by a potential concomitant CIII operation, oxidizing UQH_2_ to UQ; indeed, as shown in figure 4C, the presence of the CIII inhibitor myxothiazol led to a decrease in UQ in the combined presence of CI inhibitors and atpenin A5, implying diminished availability of UQ from CIII to CI and CII; similarly, the same can be observed in NADH levels (figure 4F), meaning that UQ arising from CIII would be used by CI and CII. Collectively, the data suggest that in acute anoxia, CII operates in the direction of UQH_2_ formation, thus CI and CII compete for the same UQ pool. However, these data do not argue that CII operates *exclusively* in forward mode.

**Figure 4:**
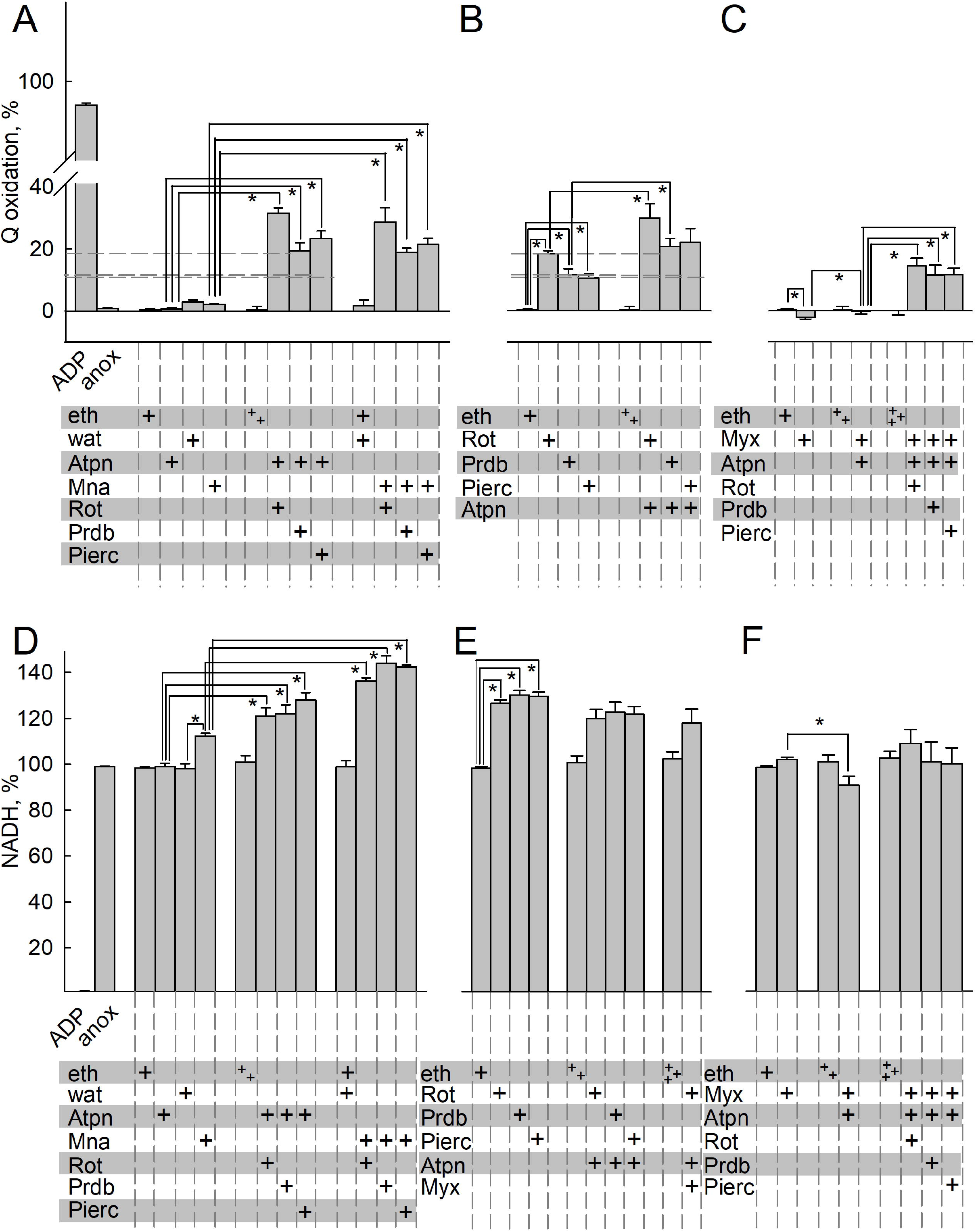
CII competes with CI for the same UQ pool during acute anoxia. Quantification of inhibitor-induced changes in UQ (A, B, C) or NADH (D, E, F) signal. Wat: water; Mna: malonate; Atpn: atpenin A5. The additions of the compounds mentioned in the bottom of each panel were made from top to bottom. Data pooled from 3-37 independent experiments. * p<0.05. Whenever two or three crosses are indicated within a single box in the bottom of a panel mentioning additions of chemicals, they signify double or triple addition of the vehicle.

### CII operates mostly towards UQH_2_ formation, but also in reverse, reducing fumarate during acute anoxia

To address the directionality of CII during acute anoxia, we examined the effect of metabolites passing through CII by measuring UQ oxidation % and NADH %. As shown in figure 5A, addition of succinate after rotenone leads to UQ reduction under anoxia, implying that CII can still operate in forward mode. In figure 5B, addition of succinate before rotenone in anoxic mitochondria depicts lesser inhibition of UQH_2_ formation than rotenone alone (figure 5A), implying that succinate was impairing CI operation during anoxia. This effect of succinate was abolished by atpenin, shown in figure 5C. Addition of any CI inhibitor after succinate during anoxia led to an increase in NADH levels implying that CI was still operational (figure 5F). However, the increase in NADH levels is smaller in the presence of exogenously added succinate than in its absence (compare dash line connecting figure 5E with 5F). Likewise, atpenin A5 abolished the effect of succinate on any CI inhibitor-induced changes on NADH levels (see figure 5G). These data reiterate that CII operates in forward mode, and this hinders CI’s ability to produce NADH, most likely because CII ‘steals’ from the finite pool of UQ available during acute anoxia, from CI. It remains unexplained, however, why exogenous addition of succinate in the presence of rotenone still yields a decrease in NADH levels.

**Figure 5:**
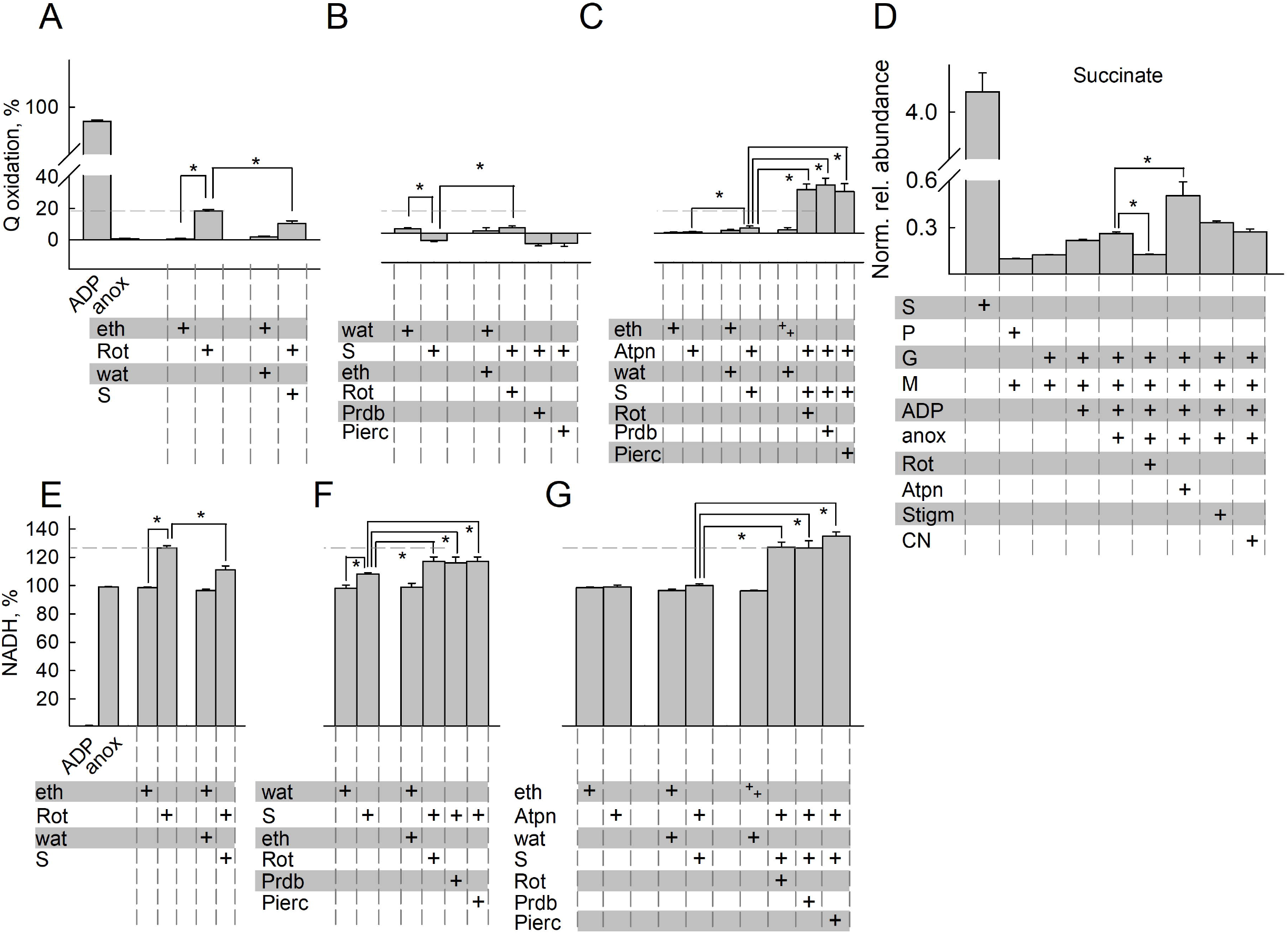
CII operates mostly towards UQH_2_ formation, but also in reverse, reducing fumarate during acute anoxia. Quantification of inhibitor-induced changes in UQ (A, B, C) or NAD (E, F, G) signal. D: Untargeted metabolomic analysis of succinate (S) present in the pellets of mitochondria treated with the conditions indicated in the panel. The y-axis illustrates normalized, log transformed, and scaled peak area. The additions of the compounds mentioned in the bottom of each panel were made from top to bottom. Arsn: arsenite; G: glutamate; M: malate; P: pyruvate; n.d.: not detected. Data pooled from 3-37 independent experiments. * p<0.05. Whenever two crosses are indicated within a single box in the bottom of a panel mentioning additions of chemicals, they signify double addition of the vehicle).

Next, to corroborate the finding that during acute anoxia CII operates in the direction towards succinate oxidation forming UQH_2_, we performed untargeted metabolomic analysis of pertinent metabolites while ETS components where pharmacologically inhibited. Mitochondria were allowed to respire on glutamate and malate in the presence of ADP to the point of O_2_ depletion from their suspending medium. This was verified by monitoring O_2_ concentration polarographically. As shown in figure 5D, anoxia increases the concentration of succinate, while rotenone partially abolishes this. Quantification of other untargeted metabolites is shown in the supplementary figure 4. On the other hand, atpenin A5 not only fails to diminish succinate concentration, it even potentiates the increase. This means that during anoxia i) CII was operating in the direction of succinate oxidation (i.e. forward mode) and that ii) CI operation was supporting succinate formation.

### Oxidation of NADH by CI supports OgDHC, in turn maintaining the oxidative decarboxylation pathway from glutamate

To address the possibility that CI was favoring the formation of succinate through supporting the oxidative decarboxylation of 2-oxoglutarate, we performed the untargeted metabolomic analysis in the presence of arsenite (HAsO_3_^2-^). When using glutamate (plus malate) as a substrate, the only target of arsenite is OgDHC, if the variable to be measured is succinate concentration; glutamate dehydrogenase is not sensitive to arsenite (Papa et al., 1967). Indeed, as shown in figure 6A, arsenite abolished the increase in succinate, irrespective of the presence of rotenone or atpenin A5. This means that succinate originated from the canonical Krebs pathway: glutamate -> 2-oxoglutarate -> succinyl-CoA -> succinate. However, it does not mean that during anoxia, succinate originated *exclusively* from this metabolic branch; to address this, we traced metabolites harboring ^13^C that could originate from [U-^13^C]glutamate or [U-^13^C]malate. As shown in figure 6B, during anoxia the abundance of labelled succinate in mitochondria that were exogenously given [U-^13^C]glutamate is lower when rotenone was present; atpenin A5 increased the amount of labelled succinate, while arsenite decreased the abundance of succinate labelling, compared to its absence. The data obtained from metabolomic analysis using [U-^13^C]glutamate agree with the untargeted metabolomic analysis arguing that succinate originated from the canonical Krebs pathway, depicted in figure 6C. It is also noteworthy that in the presence of atpenin A5, while succinate is building up significantly, it is unable to exit mitochondria as easily as under other conditions (ratio of pellet to supernatant is significantly altered). The reason(s) behind this could be due to enhanced malate uptake or excretion of another dicarboxylate, effectively out-competing succinate; alternatively, this could be a result of issues co-transporting inorganic phosphate (P*i*·): the dicarboxylate transporter can exchange a dicarboxylate for P*i*· (Johnson and Chappell, 1973), so when P*i*· builds up in the matrix (due to extensive ATP hydrolysis by a reverse-operating F_1_F_O_-ATPase (Kiss *et al*., 2014)) the exchange for succinate could be inhibited, hence malate enters against citrate and the dicarboxylate carrier is effectively fully inhibited. The abundance of other targeted metabolites is shown in supplementary figure 5. There, the ~5-fold increase of oxoglutarate from [U-^13^C]glutamate by arsenite indicates genuine inhibition of the oxoglutarate complex. Results on labeled fumarate also confirm that while fumarate hydratase is working at equilibrium loading the pool from unlabeled malate, contribution from glutamate is significantly reduced after rotenone, atpenin A5 and arsenite. The reduction after rotenone is highly similar to that of succinate, showing the reduction is upstream, as opposed to the presence of atpenin A5. However, when using [U-^13^C]malate, there was succinate labelling during anoxia (figure 6D) but with no labelling of oxoglutarate (figure 6E). This could only mean that CII was also operating in reverse to some extent, depicted in figure 6F, and consistent with the notion that most of the citrate being produced is being exported against the malate gradient. The abundance of other metabolites that acquired labelling from [U-^13^C]malate is shown in supplementary figure 5. Regarding schemes 6C and 6F even though the reaction catalyzing the interconversion of malate to oxaloacetate by malate dehydrogenase is definitely occurring as evidenced by the aspartate labeling, oxaloacetate is omitted as we could not detect it. In schemes 6C and 6F we only mark metabolites outlining the metabolic routes taken, and not all metabolites detected with the ^13^C labels, for clarity. So how can it be that CII operates mostly in forward but also in reverse, under the same circumstances? The reaction catalyzed by CII is succinate + UQ <-> fumarate + UQH_2_; however, there are other reactions affecting each reactant as well: succinate is also influenced by succinyl-CoA ligase, fumarate by fumarase, UQ and UQH_2_ by CI and other enzymes converging at the mitochondrial Q junction (Banerjee et al., 2021). By plotting succinate/fumarate ratio (on the basis of metabolite abundance in the pellets during anoxia, obtained from the targeted metabolomic analysis) *vs* UQ/UQH_2_ ratio, it is apparent that CII directionality is dictated by the pair of values across a straight line, see figure 6G. However, fumarase, CI, and enzymes converging at the mitochondrial Q junction (as well as other enzymes having succinate or fumarate as substrates or products, see https://metabolicatlas.org/explore/Mouse-GEM/gem-browser/metabolite/MAM02943m and https://metabolicatlas.org/explore/Mouse-GEM/gem-browser/metabolite/MAM01862m, respectively) will lead to a shifting pattern of succinate/fumarate and UQ/UQH_2_ pair values across this line, which means that the direction favored by CII will be changing as dictated by the very same value pairs. In addition, since UQ may be protein-bound or unbound to the extent of 10-32% (Lass and Sohal, 1999) the amount of protein-unbound UQ and the parameters influencing this binding (such as the lipophilicity property of the UQ type (Briere et al., 2004), (Chen et al., 1986), (Lenaz, 1998)) will also dictate CII directionality.

**Figure 6:**
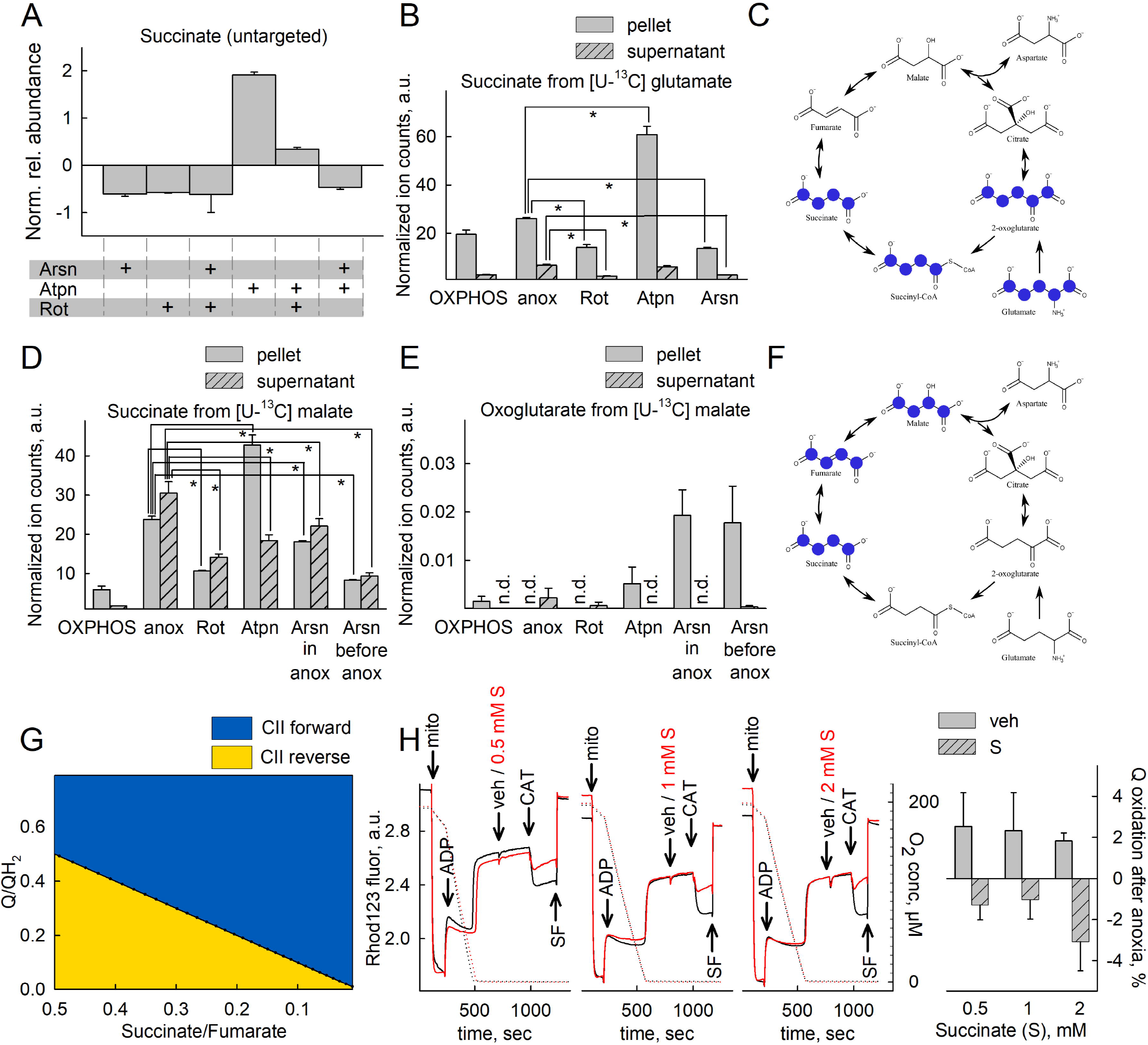
Oxidation of NADH by CI supports OgDHC, in turn maintaining the oxidative decarboxylation of 2-oxoglutarate during acute anoxia. A: Untargeted metabolomic analysis of succinate present in the pellets of mitochondria treated with the conditions indicated in the panel. The y-axis illustrates normalized, log transformed, and scaled peak area. B, D: Targeted metabolomic analysis of succinate present in the effluxes (supernatants) or pellets of mitochondria treated with the conditions indicated in the panel, when using [U-^13^C]glutamate (panel B) or [U-^13^C]malate (panel D). In E, labelling in oxoglutarate from [U-^13^C]malate is shown. C, F: Schemes illustrating the paths of ^13^C labels in glutamate or malate to succinate during anoxia, respectively. Marvin was used for drawing chemical structures, Marvin version 21.18, ChemAxon, (https://www.chemaxon.com). G: 2D plot of CII directionality depicted from succinate/fumarate ratio as a function of Q/QH_2_ ratio. H: Succinate (0.5, 1, 2 mM) diminishes the CAT-induced changes in rhodamine 123 fluorescence (solid lines, Rhod123, arbitrary units, three leftmost subpanels) in anoxic mouse liver mitochondria (oxygen concentration in μM, shown in dotted lines) in addition to reducing UQ to UQH_2_ (rightmost subpanel). Data pooled from 3-6 independent experiments. * p<0.05.

Mindful that succinate may arise from either i) glutamate -> 2-oxoglutarate -> succinyl-CoA -> succinate implying CII forward mode (since the succinyl-CoA to succinate is catalyzed by the fully reversible succinyl-CoA ligase thus, succinate removal is imperative (Chinopoulos *et al*., 2010)), and/or ii) reduction of fumarate implying CII reversal, the question arises which pathway contributes more significantly to altering succinate concentration in the matrix during acute anoxia. To this end, we titrated the amount of exogenously added succinate in reversing ANT directionality during anoxia; the rationale of this approach is that if CII reversal is considerable, the amount of succinate generated should inhibit the ATP-forming mtSLP which dictates ANT directionality (Chinopoulos *et al*., 2010). As shown in figure 6H the amount of exogenously added succinate for causing CAT-induced delayed loss of Δ*Ψ*_mt_ after anoxia is in the order of 0.5-2 mM (left three panels). The K_*m*_ of the dicarboxylate transporter translocating succinate across the inner mitochondrial membrane is > 1.1 mM (Palmieri et al., 1971). Accumulation of succinate in the matrix is also dependent on Δ*Ψ*_mt_; the more depolarized mitochondria are, the less succinate is taken up (Quagliariello and Palmieri, 1968). It is therefore reasonable to assume that addition of 0.5 mM succinate to already depolarized mitochondria due to anoxia leads to a small increase in matrix succinate concentration, well below 0.5 mM. As this is a small amount, we conclude that CII could not operate in reverse to an appreciable degree during acute anoxia, as the amount of succinate formed is insufficient to hinder mtSLP. Furthermore, exogenously added 0.5 mM succinate still yielded UQ reduction to UQH_2_, implying that CII was mostly operating in forward mode (figure 6H, rightmost panel). Although we cannot quantify the extent of contribution of CII reversal in yielding succinate, we conclude that this path is quantitatively much smaller than the canonical route.

## Materials and Methods

### Animals

Mice were of C57Bl/6 background. The animals used in our study were of either sex and between 2 and 6 months of age. Mice were housed in a room maintained at 20–22 °C on a 12-h light–dark cycle with food and water available ad libitum, unless otherwise indicated. The study was conducted according to the guidelines of the Declaration of Helsinki, and were approved by the Animal Care and Use Committee of the Semmelweis University *(Egyetemi Állatkísérleti Bizottság*, protocol code F16-00177 [A5753-01]; date of approval: May 15, 2017).

### Mitochondrial isolation

Liver mitochondria were isolated from mice as described in (Chinopoulos *et al*., 2009). Protein concentration was determined using the bicinchoninic acid assay, and calibrated using bovine serum standards (Smith et al., 1985) using a Tecan Infinite^®^ 200 PRO series plate reader (Tecan Deutschland GmbH, Crailsheim, Germany).

### Determination of membrane potential (ΔΨ_mt_) in isolated mitochondria

Δ*Ψ*_mt_ of isolated mitochondria (0.5-1 mg mouse liver mitochondria in 2 ml buffer medium, the composition of which is described in (Chinopoulos *et al*., 2010)) was estimated fluorimetrically with rhodamine 123 (Emaus et al., 1986). Fluorescence was recorded using the NextGen-O_2_k prototype equipped with the O_2_k-Fluo Smart Module, with optical sensors including a LED (465 nm; <505 nm short-pass excitation filter), a photodiode and specific optical filters (>560 nm long-pass emission filter) (Krumschnabel et al., 2014). Experiments were performed at 37 °C.

### Mitochondrial respiration

Oxygen consumption was monitored polarographically using an Oxygraph-2k. 0.5-1 mg of mouse liver mitochondria were suspended in 2 ml incubation medium, the composition of which was identical to that for Δ*Ψ*_mt_ determination. Experiments were performed at 37 °C. Oxygen concentration (μM) and oxygen flux (pmol·s ^-1^·mg ^-1^; negative time derivative of oxygen concentration, divided by mitochondrial mass per volume and corrected for instrumental background oxygen flux arising from oxygen consumption of the oxygen sensor and back-diffusion into the chamber) were recorded using DatLab software (Oroboros Instruments).

### Determination of NADH autofluorescence in isolated mitochondria

NADH autofluorescence was measured using the NADH-Module of the NextGen-O_2_k (Oroboros Instruments). The NextGen-O_2_k allows simultaneous measurement of oxygen consumption and NADH autofluorescence, incorporating an ultraviolet (UV) LED with an excitation wavelength of 365 nm and an integrated spectrometer which records a wavelength range between 450 and 590 nm. The light intensity of the LED was set to 10 mA. 0.5-1 mg of mouse liver mitochondria were suspended in 2 ml incubation medium, the composition of which was identical to that for Δ*Ψ*_mt_ determination, as described in (Chinopoulos *et al*., 2010). Experiments were performed at 37 °C.

### Mitochondrial UQ redox state

Coenzyme UQ redox state of isolated mitochondria suspended in a buffer composition as described in (Chinopoulos *et al*., 2010) was followed amperometically using a three electrode system with coenzyme Q_2_ (CoQ_2_, 1 μM) as mediator, using the Q-Module of the NextGen-O_2_k (Komlódi T, 2021). The reference electrode was Ag/AgCl/(3M KCl). The auxiliary electrode was made of platinum and the working electrode was fabricated from glassy carbon. Oxidation peak potential of CoQ_2_ measured by cyclic voltammetry was set to the glassy carbon to measure the oxidation of reduced CoQ_2_. UQ redox state was recorded simultaneously with O_2_ flux and rhodamine 123 fluorescence. All drugs used in this study were verified not to exert any artefactual alterations on the electrodes of Q module using cyclic voltammetry, shown in supplementary figure 6.

### Mitochondrial swelling

Swelling of isolated mitochondria was assessed by measuring light scatter at 520 nm (37 °C) in a Hitachi F-7000 fluorescence spectrophotometer. 0.5 mg of mouse liver mitochondria were suspended in 2 ml incubation medium, the composition of which was identical to that for Δ*Ψ*_mt_ determination, as described in (Chinopoulos *et al*., 2010). Experiments were performed at 37 °C. Anoxic conditions were achieved by manufacturing a custom-made plug for polymethacrylate cuvettes by 3D-printing. The plug 3D design and instructions for use are published in https://www.thingiverse.com/thing:3156148. At the end of each experiment, the non-selective pore-forming peptide alamethicin (80 μg) was added as a calibration standard to cause maximal swelling.

### Prothrombin time measurement

Blood samples were taken from the saphenous vein in 110 mM Na3-citrate (citrate/blood volume ratio 1:10), as described in (Parasuraman et al., 2010). After centrifugation at 2,500*g* for 15 min the plasma supernatant was collected and used for the measurement within 4 hours. Prothrombin time was measured with Technoplastin-HIS reagent (Technoclone Herstellung von Diagnostika und Arzneimitteln GmbH, Vienna, Austria) in a coagulometer KC-1A (Amelung, Lemgo, Germany) as the time to form clots from 100 μl plasma diluted 2-fold in 10 mM HEPES 150 mM NaCl pH 7.4 buffer by 200 μl Technoplastin-HIS reagent.

### Untargeted metabolomic analysis

Mitochondrial suspensions were ‘spiked’ with 1 mM L-norleucine, a non-metabolizable substrate that yields a highly recognizable signature during the metabolite analysis and was used for normalizing volumes and keeping pipetting errors at check. At specified times during the experiments (indicated in the text) three 0.6 ml aliquots from the 2 ml mitochondrial suspensions were spun at 14,000 rpm for 5 min at 4 °C. Supernatants (0.5 ml) were separated from the pellets. Any remaining visible aqueous phase was removed from the pellets and discarded. Both pellets and supernatants were transferred to new tubes and to each tube 0.5 ml of ice-cold 80% methanol was added. The tubes were snap frozen in liquid nitrogen during processing to ensure samples were kept cold and transferred to −80 °C until further processing. Approximately two days later, samples were sonicated for 10 min at room temperature and then centrifuged at 14,000 rpm for 10 min at 4 °C. All of the supernatants were removed, transferred to Eppendorf tubes and evaporated to dryness overnight using a centrifugal evaporator. Once dry, the dried lysates were stored at −80 °C until further analysis. Dried lysates were reconstituted in 2:1:1 acetonitrile:MeOH:H_2_O, to yield a concentration of 200 mg/ml and spun at 14,000 rpm for 10 min at 4 °C to remove excess debris before analysis. Chromatography was performed using an Agilent 1290 Infinity UPLC. Ten microliters of each sample were injected onto a ZIC-pHILIC column (EMD Millipore, Billerica, MA) with dimensions of 150 × 4.6 mm, 5 μm. Metabolites were separated using an acetonitrile/H_2_O with 20 mM ammonium carbonate (pH 9.2) gradient over a 29-min period. A 10-min re-equilibration time was carried out in between injections. Detection was performed using an Agilent 6550 quadrupole-time-of-flight (QToF) mass spectrometer, operated in both negative and positive modes. Full scan MS data was collected from m/z 70–1000 and metabolites were identified in an untargeted manner by looking within 10 ppm of the expected m/z values. Real-time mass calibration was performed throughout the duration of sample analysis. Data was processed using a publically available software package, MAVEN (Clasquin et al., 2012). Area under the chromatographic peak for each metabolite was calculated and exported to assess for differences in metabolite abundances.

### Targeted metabolomic analysis

[U-^13^C]glutamate (5 mM) or [U-^13^C]malate (2.5 mM) was added to the mitochondrial suspensions, as indicated in the text. Metabolites extraction was performed similar to the untargeted metabolite analysis, with the exception that in the supernatants, 1 mM methionine was added for normalizing volumes, as it also yields a highly recognizable signature during the analysis. Dried lysates were derivatized using a two-step protocol. Samples were first treated with 2% methoxamine in pyridine (40 μl, 1 h at 60°C), followed by addition of N-(tert-butyldimethylsilyl)-N-methyl-trifluoroacetamide, with 1% tert-butyldimethylchlorosilan (50 μl, 1 h at 60°C). Samples were transferred to glass vials for GC-MS analysis using an Agilent 8890 GC and 5977B MSD system. 1 μL of sample was injected in splitless mode with helium carrier gas at a rate of 1.0 mL·min-^1^. Initial GC oven temperature was held at 100°C for 1 minute before ramping to 160°C at a rate of 10°C·min-^1^, followed by a ramp to 200°C at a rate of 5°C·min-^1^ and a final ramp to 320°C at a rate of 10°C·min-^1^ with a 5 minute hold. Compound detection was carried out in scan mode. Total ion counts of each metabolite were normalized to the internal standard norleucine (supernatants) or methionine (pellets).

### Quinones extraction

2 mg mitochondria were suspended in a solution of 5 mM ferricyanide, 100 mM Tris, pH 8.0. Proteins were precipitated by addition of methanol triple the aqueous volume, followed by extraction three times with light petroleum (bp. 40-60° C). After evaporation of the solvent from the extract, the residues were dissolved in ethanol and the absorption measured at 275 nm before and after stepwise additions of a 5 g/l borohydride solution in a 1 cm path length cuvette. Quinone concentration was calculated with the oxidized-reduced difference absorbance Δε_275 nm_ = 12.5 mM^-1^·cm^-1^ (Crane and Barr, 1971).

### Reagents

Standard laboratory chemicals were from Sigma Aldrich (St Louis, Missouri, US). SF6847 and atpenin A5 were purchased from Enzo Life Sciences (ELS AG, Lausen, Switzerland). Mitochondrial substrates were dissolved in bi-distilled water and titrated to pH 7.0 with KOH. ADP was purchased as a K^+^ salt of the highest purity available (Merck) and titrated to pH 6.9. Concentrations of glutamate (G), malate (M), succinate (S) and oxoglutarate (Og) were always 5 mM when present. ADP concentrations were 2 mM. Rotenone (Rot, 1 μM), myxothiazol (Myx, 0.1 μM), stigmatellin (Stigm, 0.5 μM) carboxyatractyloside (CAT, 1 μM), SF6847 (SF, 0.25 μM). Both myxothiazol and stigmatellin block CIII; they have been used in our experiments according to availability.

### Statistics

Data are presented as averages ± SEM. Significant differences between two groups were evaluated by Student’s t-test or Mann-Whitney U Test if normality failed. Significant differences between three or more groups were evaluated by one-way ANOVA or ANOVA on Ranks if normality failed; statistical significance was accepted when p < 0.05 (shown as *).

## Discussion

The catabolism of glutamine by the oxidative decarboxylation route entering the Krebs cycle during hypoxia has been firmly established (Zhang *et al*., 2018), (Kohlhauer et al., 2018), (Kiss *et al*., 2014), (Seyfried *et al*., 2020). What has not been fully addressed is the origin of NAD^+^ for supporting OgDHC in the pathway glutamine -> glutamate -> 2-oxoglutarate -> succinyl-CoA -> succinate, for the 2-oxoglutarate to succinyl-CoA conversion, mindful of the highly reductive matrix environment during hypoxia. To this end, various potential sources have been theorized to contribute (Chinopoulos, 2020). The most important finding of the present study is that during anoxia CI remains operational, albeit to a diminished, but sufficient extent for supporting the reaction catalyzed by OgDHC with NAD^+^.

The significance of this finding has the following four ramifications:

i. it can explain earlier findings reporting that CI inhibitors yield less succinate when added in hypoxic tissues (Hoberman and Prosky, 1967), (Hohl et al., 1987), (Zhang *et al*., 2018). The accumulation of succinate in ischemia is unquestionable (Chouchani et al., 2014), (Hochachka and Dressendorfer, 1976), (Hochachka et al., 1975), (Tretter et al., 2016), however, its origin is debated; on one hand, CII reversal reducing fumarate to succinate yielding UQ for CI has been convincingly demonstrated (Chouchani *et al*., 2014), (Spinelli *et al*., 2021); on the other, succinate formation by the oxidative deamination of glutamine/glutamate and following canonical Krebs cycle activity has been unequivocally shown to be quantitatively more important (Zhang *et al*., 2018). Here we show that in the presence of sufficient UQ stores and for as long as they may last during acute anoxia, the canonical Krebs cycle activity is the most prominent path for succinate formation; in either case -CII reversal yielding succinate from fumarate or CII forward diverting UQ away from CI-CI remains operational in anoxia, providing OgDHC with NAD^+^. The ability of the tissue to withstand anoxia will eventually depend on the availability of UQ pools for the acute phase and the availability of fumarate for the latent phase. Fumarate supporting CI and in extension of this both Δ*Ψ*_mt_ (Pell et al., 2016) and provision of NAD^+^ to OgDHC (this study) could arise from aspartate through the concerted action of purine nucleotide cycle and aspartate aminotransferase (Chouchani *et al*., 2014). However, there are two concerns with this concept: first, the purine nucleotide cycle is an energy-dependent process (Idstrom et al., 1990); second, the energy provided by CI could not be in the form of Δ*Ψ*_mt_ to a sufficient extent: acknowledging that CI activity in anoxia is ~10% of its theoretical maximum and mindful that only 4 protons are pumped out of the 4+4+2 for CI, CIII and CIV, respectively by the ETS and that Δ*Ψ*_mt_ fluctuates between-108 and −158 mV in normoxia (Gerencser et al., 2012) and ~-100 mV in anoxia (Kiss *et al*., 2014), the contribution of CI to generating membrane potential in anoxia should be a meager 2.5 - 3.7 mV, which is reflected in our rhodamine 123 recordings, given the evanescent changes in fluorescence. Thus, it is likely that the availability of UQ for CI for the acute phase shown hereby may generate sufficient UQH_2_ that can support CII reversal in the latent phase.
ii. Oxidation of UQH_2_ in mitochondria is necessary for tumor growth (Martinez-Reyes et al., 2020), where oxygen availability is frequently limited (Vaupel and Harrison, 2004): it is extremely likely that NADH oxidation by CI requiring UQ in hypoxia is the missing link for providing NAD^+^ to OgDHC supporting the oxidative decarboxylation branch of glutaminolysis; glutaminolysis is hallmark of many cancers (Wise and Thompson, 2010). Indeed, the group of Chandel (Martinez-Reyes *et al*.,2020) demonstrated that the loss of CI thwarted tumor growth and this was rescued by mitochondrially-targeted expression of the NADH oxidase LbNOX (Titov et al., 2016).
iii. Relevant to the results published in (Martinez-Reyes *et al*., 2020), the role of CI in the proliferation of cancer cells has been reviewed in (Urra et al., 2017), and the consensus was that a complete-as opposed to an incomplete-loss of CI activity halts tumor growth, although avoidance of reactive oxygen species (ROS) generation was proposed to mediate this effect. In addition, the CI inhibitor EVT-701 exhibiting potency against solid cancers in murine models and human cell lines is in line for being evaluated in clinical trials (Luna Yolba et al., 2021).
iv. CI has been described to exist in two forms, an active (*A*) and a de-active (*D*) form, the latter signifying a dormant state of the complex which is not inactivated or covalently modified in any way (Babot et al., 2014); the *A* to *D* transition occurs during ischemia, *i.e*. when substrate and oxygen availability are limited (Babot and Galkin, 2013). The *D* form is thought to exert a protective role by delaying the rate of regaining normal respiration rates during reoxygenation, thus potentially mitigating ROS production (Stepanova et al., 2019); the other side of the coin is that when in *D* form, CI risks becoming permanently inactive (Galkin et al., 2009). Although it is not possible to decipher if residual CI activity maintained by UQ availability during acute anoxia will be beneficial or not regarding ROS formation *vs* risk of permanent inactivation, it certainly influences the *A* to *D* transition phenomenon.

A common denominator of the above is that subsequent catabolism of succinyl-CoA, the product of OgDHC which is supported with NAD^+^ provided by residual CI activity during anoxia will contribute to mtSLP. The generation of high-energy nucleotides by mtSLP has been unjustly side-lined in the literature; it is true that the rate of ATP (or GTP, depending on subunit composition of the succinyl-CoA ligase and the presence of a nucleoside diphosphokinase (Lambeth et al., 2004), (Kacso et al., 2016)) synthesis dwarfs compared to that by the mitochondrial F1FO-ATPase. However, during anoxia the F_1_F_O_-ATPase is hydrolyzing ATP instead of producing it (St-Pierre et al., 2000); in this case, the amount of ATP produced by mtSLP in combination to the much smaller volume of mitochondrial network compared to that of the cytosol amplifies the effects of ATP production in terms of concentration. This could rescue a tissue experiencing respiratory inhibition by preventing mitochondria from becoming ATP drains (Chinopoulos, 2011b).

In conclusion, we hereby showed that in isolated mouse liver mitochondria experiencing anoxia, CI exhibits sufficient activity yielding NAD^+^ for OgDHC that in turns forms succinyl-CoA supporting mtSLP; the availability of UQ is a critical factor for this residual CI activity. Forward operation of CII is necessary for maintaining succinate levels low enough to allow the reversible succinyl-CoA ligase reaction to proceed towards ATP (or GTP) synthesis, but on the other hand, CII reversal also occurs reducing fumarate to succinate and oxidizing UQH_2_ to Q, the latter favoring CI activity. All other enzymatic reactions in which succinate, fumarate, UQ and UQH_2_ participate, contribute to CII directionality and ultimately influence CI residual activity.

## Limitation of present study

A main concept of the present work is the demonstration that UQ availability dictates CI activity; therefore, manipulation of UQ pools by disrupting UQ biosynthetic pathways could have been argued to lend strong support to the above claim. However, disrupting UQ biosynthesis leads to impaired mitochondria (Hargreaves et al., 2020); it would be exceedingly difficult to achieve anoxic state in impaired mitochondria due to negligible OXPHOS state of respiration. Thus, we have alternatively relied on limiting vitamin K supplementation to mice; “vitamin K” is actually a group of naphthoquinones that includes menaquinone and phylloquinone (Shearer and Okano, 2018). These two quinones carry electrons in bacteria and plants, respectively, whereas eukaryotes use ubiquinone (UQ). However, the eukaryotic enzyme ferroptosis suppressor protein 1 (FSP1), a NAD(P)H-ubiquinone reductase reduces both vitamin K quinones and UQ (Mishima et al., 2022) and by doing this, vitamin K may “spare” UQ pools. Since vitamin K is a regular dietary constituent, we reasoned that by decreasing vitamin K availability in a relatively acute manner (mice were subject to a vitamin K-deficient diet for 1-3 weeks) UQ pools could be sufficiently diminished but not to the extent of impairing OXPHOS respiration.

## Acknowledgments

This work was supported by grants from NKFIH KH129567, NKFIH K135027 and TKP2021-EGA-25 to C.C., grants NKFIH K137563 and TKP2021-EGA-24 to K.K. and from the project NextGen-O_2_k (Oroboros Instruments) which has received funding from the European Union’s Horizon 2020 research and innovation programme under grant agreement N^<u>o</u>^ 859770. We would like to acknowledge the support and resources of the Birmingham Metabolic Tracer Analysis Core. T.N.S. has received funding from Childhood Cancer UK, Foundation for Metabolic Cancer Therapies, the Corkin Family Foundation, and Dr. Edward Miller. D. R. was supported by a scholarship from School of PhD Studies of Semmelweis University, project no EFOP-3.6.3-VEKOP-16-2017-00009.

## Author contributions

D.R., D.B., S.N., G.P., N.K., J.R., B.G. A.K., C.H., T.K., C.D., A.R., B.C. and C.C. conducted the experiments; J.R., A.K., M.K. and D.A.T. interpreted the metabolomics results; K.K. interpreted the hemostasis results; D.A.T., E.G., M.K., N.N.and T.N.S. edited the paper; C.C. designed the experiments and wrote the paper.

