## Supplementary material for "Residual Complex I activity supports glutamate catabolism and mtSLP via canonical Krebs cycle activity during acute anoxia without OXPHOS": Legends to supplementary figures

**Legend to supplementary figure 1**: Directionality of CI operation: forward *vs* reverse operation (the latter is represented by the shaded area) as a function of QH_2_/Q, NAD^+^/NADH, matrix pH (pH_in_), ΔΨ_mt_ and ΔpH across the inner mitochondrial membrane and highlighted area indicating plausible conditions of anoxia in isolated mouse liver mitochondria. Panels to the left share the same y-axis with those corresponding to the right. All panels share the same x-axis.

**Legend to supplementary figure 2**: Oxygen concentration (top panels, in μM) recorded simultaneously with either Q redox state (A, B, E, F, G, H, I, J, K, middle panels) or NADH autofluorescence (C, D, middle panels) and/or rhodamine 123 fluorescence (Rhod123, bottom panels) in isolated mouse liver mitochondria. For experiments shown in panels A, B, C and D, substrates were glutamate and malate (5 mM each) present in the buffer prior to addition of mitochondria. Rhodamine 123 fluorescence indicative of ΔΨ_mt_ (arbitrary units, a.u.) was recorded separately from NADH autofluorescence due to spectral overlap. Mitochondria (mito), ADP (2 mM), pyridaben (Prdb, 1 μM, panel A, C, F), or piericidin A (Pierc 1 μM, panel B, D, G), carboxyatractyloside (CAT, 1 μM, all panels), SF (SF6847, 0.25 μM, all panels), rotenone (Rot, 1 μM, panels E, H, I, J, K) were added where indicated.

**Legend to supplementary figure 3**: A, B: Light scatter (520/520 exc/em) of mouse liver mitochondria indicative of swelling; in A, experiments were identical to those performed in main figure 1D, E; the ETS inhibitor is indicated in the panel. In B, CaCl_2_ pulses (0.3 mM) is added where indicated to demonstrate the extent of swelling that these mitochondria can reveal. Substrates were glutamate and malate (5 mM each) present in the buffer prior to addition of mitochondria. Experiments on light scatter were performed using a standard cuvette-based spectrofluorimeter (Hitachi F-7000) plugged with a custom-made plug for polymethacrylate cuvettes by 3D-printing, depicted in panel C. The plug 3D design and instructions for use are published in https://www.thingiverse.com/thing:3156148.

**Legend to supplementary figure 4**: Analysis of untargeted metabolites present in the pellets of mitochondria treated with the conditions indicated in the panel. The y-axis illustrates normalized, log transformed, and scaled peak area.

**Legend to supplementary figure 5**: Targeted metabolomic analysis of metabolites present in the effluxes (supernatants) or pellets of mitochondria treated with the conditions indicated in the x-axis, when using [U-^13^C]glutamate or [U-^13^C]malate.

**Legend to supplementary figure 6**: Cyclic voltammograms in the absence (black traces) or presence (red traces) of chemicals used in this study.
