## Supplementary figures and images for "Residual Complex I activity supports glutamate catabolism and mtSLP via canonical Krebs cycle activity during acute anoxia without OXPHOS"

### Supplementary figure 1

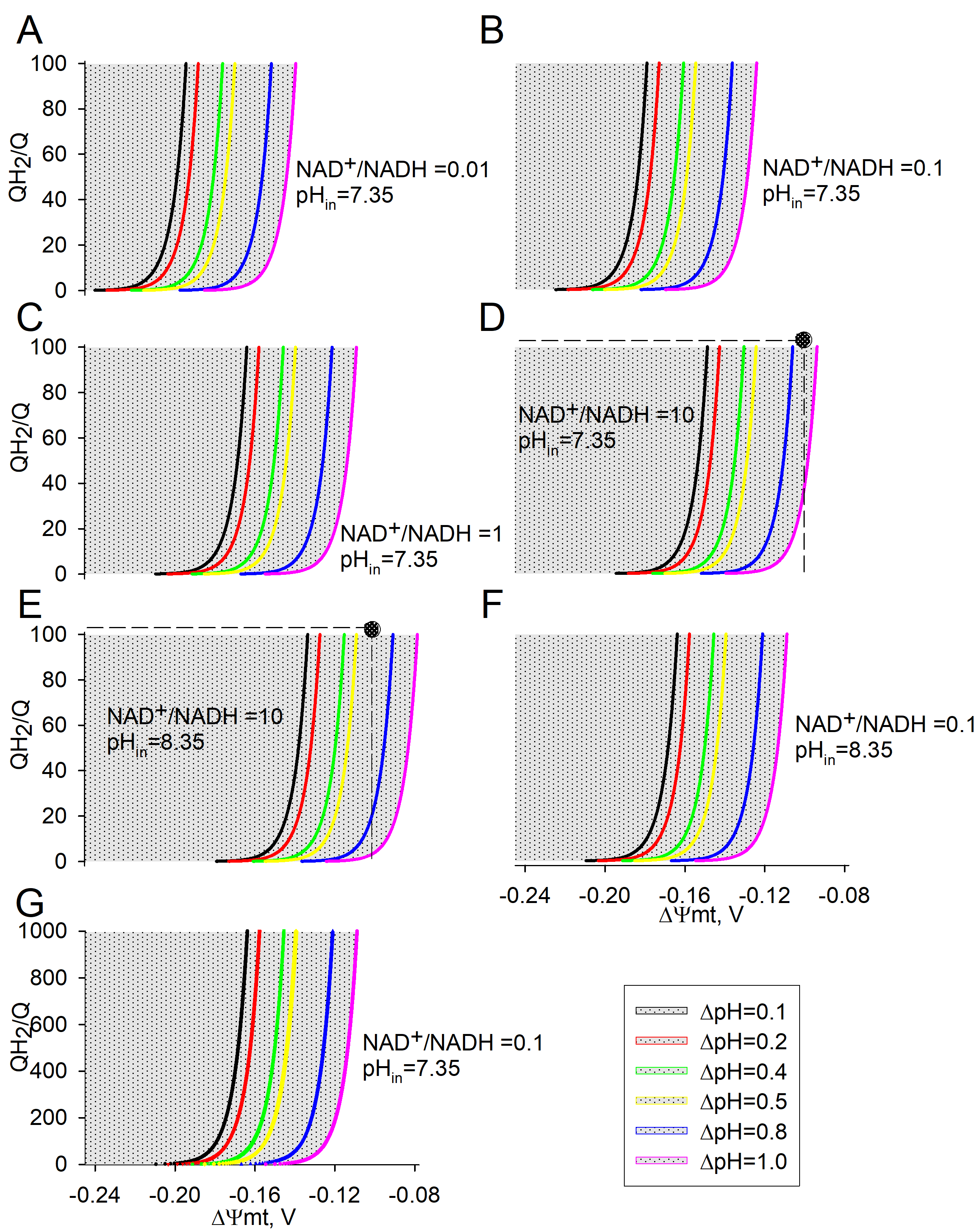

### Supplementary figure 2

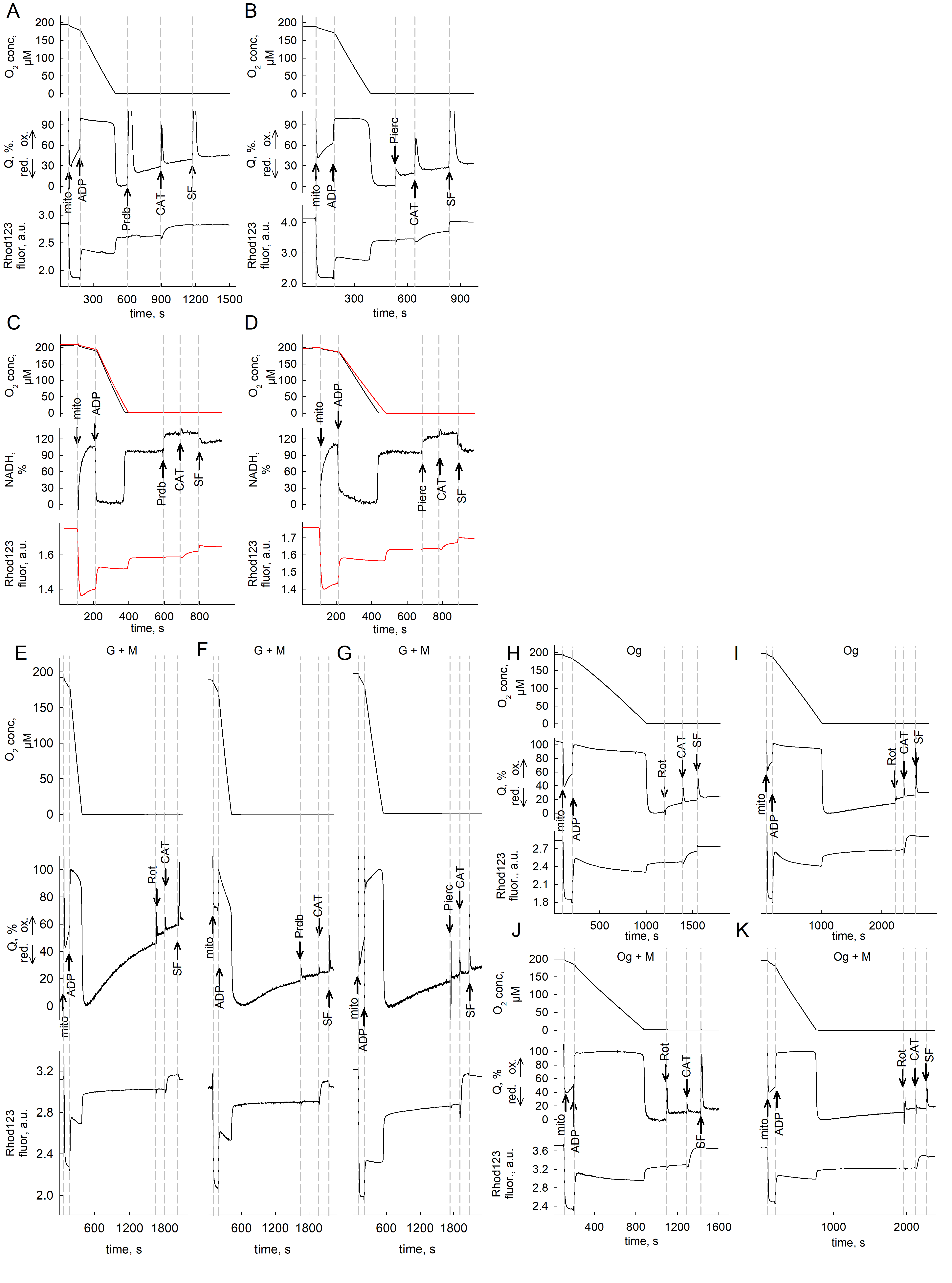

### Supplementary figure 3

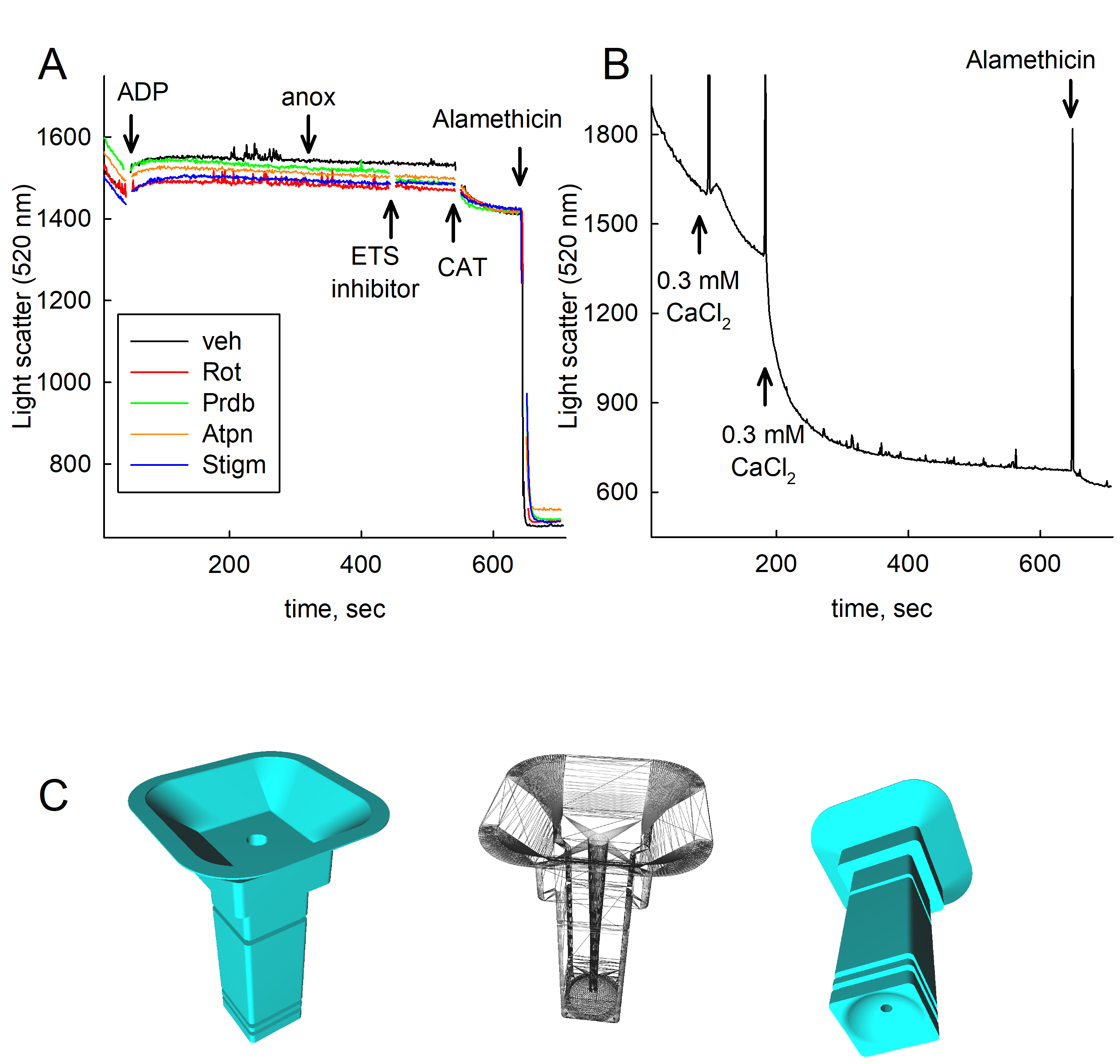

### Supplementary figure 4

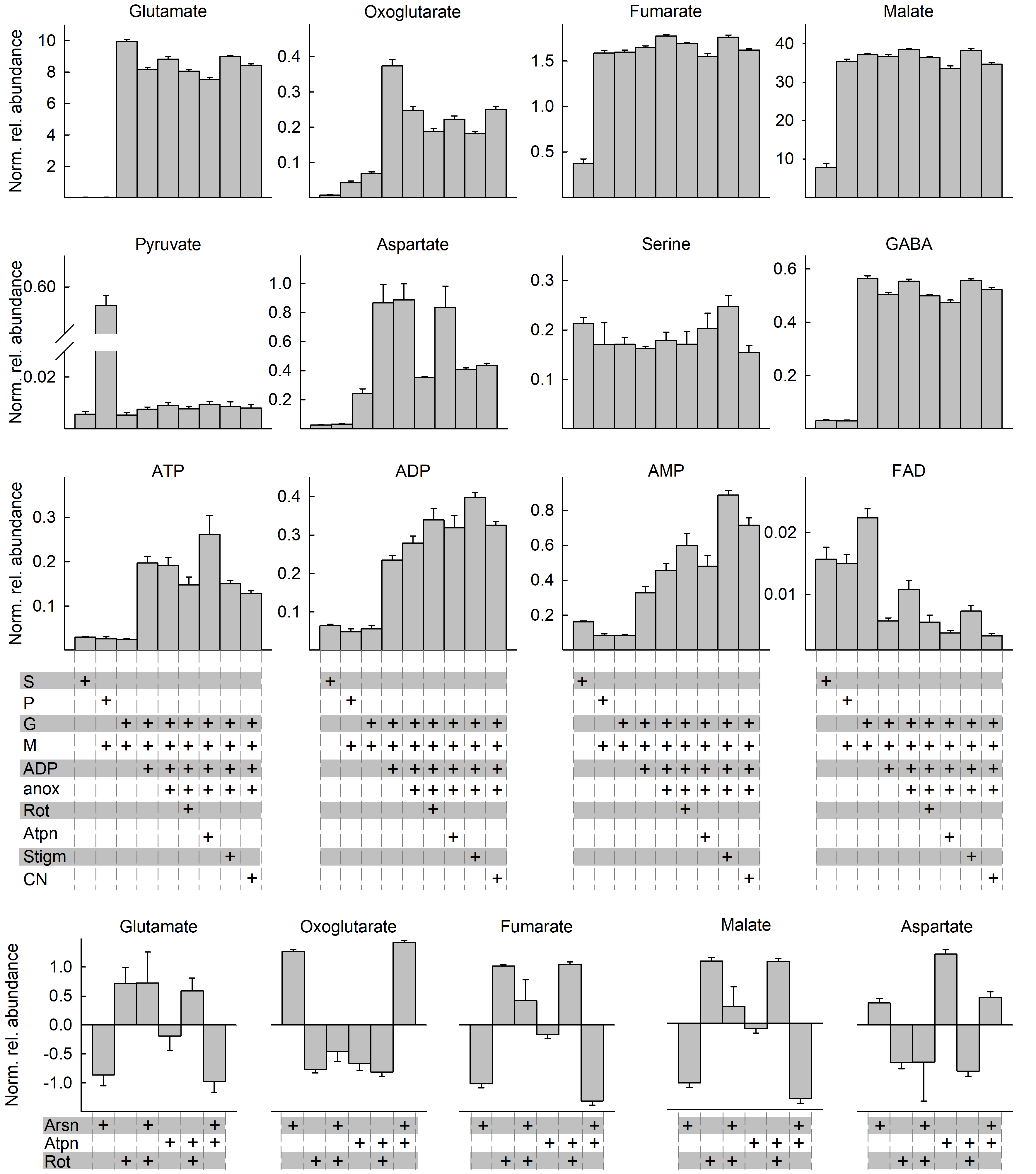

### Supplementary figure 5

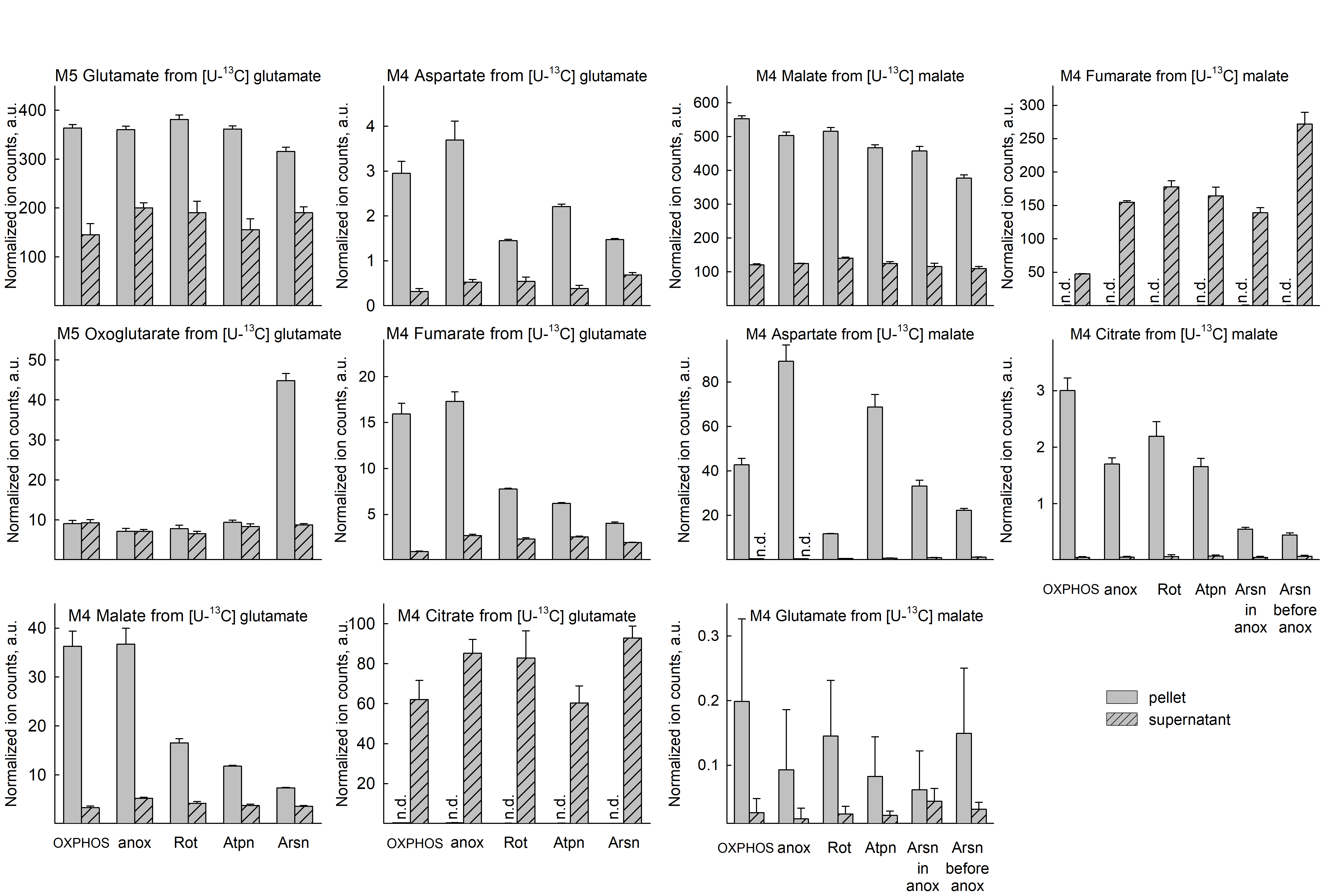

### Supplementary figure 6

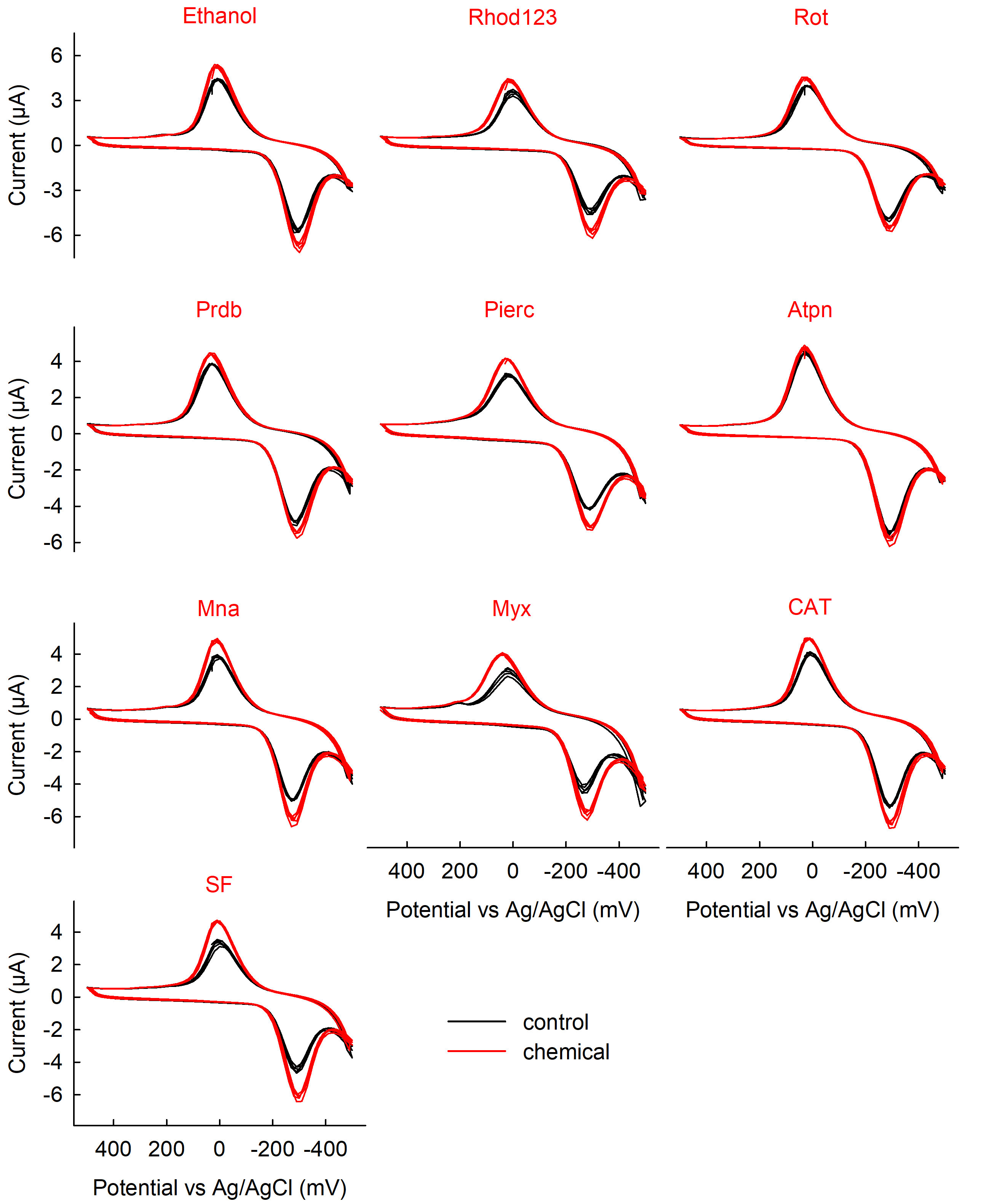
